# Distinct roles of the two SEC scaffold proteins, AFF1 and AFF4, in regulating RNA Pol II transcription elongation

**DOI:** 10.1101/2022.08.09.503397

**Authors:** Zhuanzhuan Che, Xiaoxu Liu, Qian Dai, Ke Fang, Chenghao Guo, Junjie Yue, Haitong Fang, Peng Xie, Zhuojuan Luo, Chengqi Lin

## Abstract

The P-TEFb-containing super elongation complex (SEC) plays the essential role in transcriptional elongation control. The AF4/FMR2 family members AFF1 and AFF4 are the central scaffold proteins of SEC, associated with different human diseases. However, their specific roles in transcriptional control remain unclear. Here, we report that AFF1 and AFF4 show distinct genomic distribution patterns around TSS. AFF1 mainly binds upstream of TSS, while AFF4 is enriched downstream of TSS. Pol II occupancies are reduced genome-widely after depletion of AFF1, but not AFF4. Interestingly, in a subset of active genes with strong AFF4 binding signature, AFF4 disruption causes slow elongation and early termination, while AFF1 deletion mirrors the transcriptional defects observed in the fast Pol II mutant. Furthermore, AFF4 knockdown leads to increased AFF1 levels at chromatin, and *vice versa*. In summary, our data demonstrate that AFF1 and AFF4 function, to some extent, antagonistically to ensure proper Pol II transcription.

## Introduction

In higher organisms, transcription by RNA polymerase II (Pol II) is sophisticatedly regulated at multiple stages to guarantee proper timing and output of gene expression at correct spatial positions (Conaway and Conaway, 2019; Cramer, 2019; Lis, 2019; Roeder, 2019). Recent genome wide localization studies in metazoans uncovered that subsequent to transcription initiation, Pol II is paused at the promoter-proximal regions of most genes (Jonkers and Lis, 2015; Core and Adelman, 2019). The release of paused Pol II into productive elongation by positive transcription elongation factor b (P-TEFb), consisting of cyclin dependent kinase 9 (CDK9) and CCNT1/2, is one of the rate-determining steps in transcription cycle (Marshall and Price, 1995; Wada et al., 1998; Price, 2000). It is well known that most P-TEFb is sequestered in the catalytically inactive 7SK small nuclear ribonucleoprotein particle (7SK snRNP) complex (Nguyen et al., 2001; Yang et al., 2001; Yik et al., 2003). Upon signal stimulate, P-TEFb is freed from the inactive complex, and incorporated into the active complexes, including the super elongation complex (SEC), to stimulate transcription (Bres et al., 2008; Lin et al., 2010; Lin et al., 2011).

SEC is composed of the ELL family members ELL1/2, AF9/ENL, the AF4/FMR2 family members AFF1/AFF4 and P-TEFb (Lin et al., 2010). AFF1 and AFF4 are mainly involved in the formation of SEC as the central scaffold proteins (Luo et al., 2012b). Mutations of AFF1 and AFF4 are associated with different human diseases (Smith et al., 2011; Luo et al., 2012b; Izumi et al., 2015). Previous studies have revealed that AFF1 and AFF4 can form separate SEC complexes to regulate the expression of different subsets of genes (Luo et al., 2012a; Lu et al., 2015). The AFF4 containing SEC is inclined to rapidly induce the expression of stress induced genes and many developmental genes (Lin et al., 2011). AFF1-SEC has potential to activate HIV-Tat transcription (Lu et al., 2014; Lu et al., 2015). The activation of *HSP70* upon heat shock is sensitive to the acute depletion of AFF4, but not AFF1, while the simultaneous degradation of both AFF1 and AFF4 leads to more serious effects (Luo et al., 2012a; Lu et al., 2015; Zheng et al., 2021). However, how their functional specificities are controlled remains unclear.

Recent studies have reported that abnormal elongation rate of RNA Pol II could intervene proper termination (Cortazar et al., 2019; Muniz et al., 2021). Pol II mutant with slow velocity could be chased easily by the termination factor XRN2 which cleaves the nascent RNA, and thus released from upstream of termination site; while fast elongation pushes termination further downstream (Fong et al., 2015). Loss of function studies of several elongation factors, including PAF1, SPT6, and CDK9, have shown termination defects similar to slow Pol II mutants (Ardehali et al., 2009; Booth et al., 2018; Hou et al., 2019). Treatment of KL-1/2, the small molecule inhibitors of SEC, destabilizes the proteins of AFF1, AFF4 and ELL2, disrupts the interaction of CCNT1 with AFF1 and AFF4, and leads to reduced Pol II elongation rate and defective termination (Liang et al., 2018). However, how AFF1 and AFF4 specifically function in transcriptional elongation control need to be further investigated.

Here, we examined the genome-wide occupancies of AFF1 and AFF4 in A549 cells, and found distinct distribution patterns of AFF1 and AFF4 around Pol II-bound transcription start site (TSS). AFF1 is enriched at ∼45nt upstream of TSS, while AFF4 locates at ∼175nt downstream of TSS. Depletion of AFF1 appears to have global effects on Pol II occupancies over promoters and transcribing units, while AFF4 knockdown only marginally affects Pol II occupancies in A549 cells. Interestingly, we found that in a subset of AFF4-bound active genes loss of AFF4 leads to reduced Pol II elongation rate and early termination. In contrast, depletion of AFF1 causes fast Pol II elongation and transcription readthrough. Furthermore, we also observed increased AFF4 chromatin occupancy in AFF1-depleted cells. Simultaneously, AFF1 depletion leads to increased AFF4 protein levels, and *vice versa*. Our data indicated that AFF1 and AFF4 could regulate transcription at different stages with a certain degree of antagonistic effects.

## Results

### Distinct distribution patterns of AFF1 and AFF4 around TSS

To investigate the roles of AFF1 and AFF4, we first profiled their genomic occupancy patterns in A549 cells. 13,078 AFF1 peaks and 6,555 AFF4 peaks were identified with an FDR < 0.05, respectively (Figure 1—figure supplement 1A). Majorities of AFF1 (∼65%) and AFF4 (∼74%) peaks were located at the Pol II bound transcriptional start sites (TSS) (Figure 1—figure supplement 1A and Figure 1A). Consistent with our previous reports (Lin et al., 2011), AFF4 travels with Pol II into the gene body, but mostly enriched at 5’ regions of the highly transcribed genes. Interestingly, metaplot analysis suggested that AFF1 and AFF4 peaks centered at different sites around TSS (Figure 1B-C and Figure 1—figure supplement 1B). AFF1 was co-localized with TBP at ∼45nt upstream of TSS, while AFF4 peak summits were detected at ∼175nt downstream of TSS, the binding profiles of AFF1, AFF4 and TBP at the *PLK2* gene locus were shown as an example (Figure 1B-C and Figure 1—figure supplement 1B).

**Figure 1.**
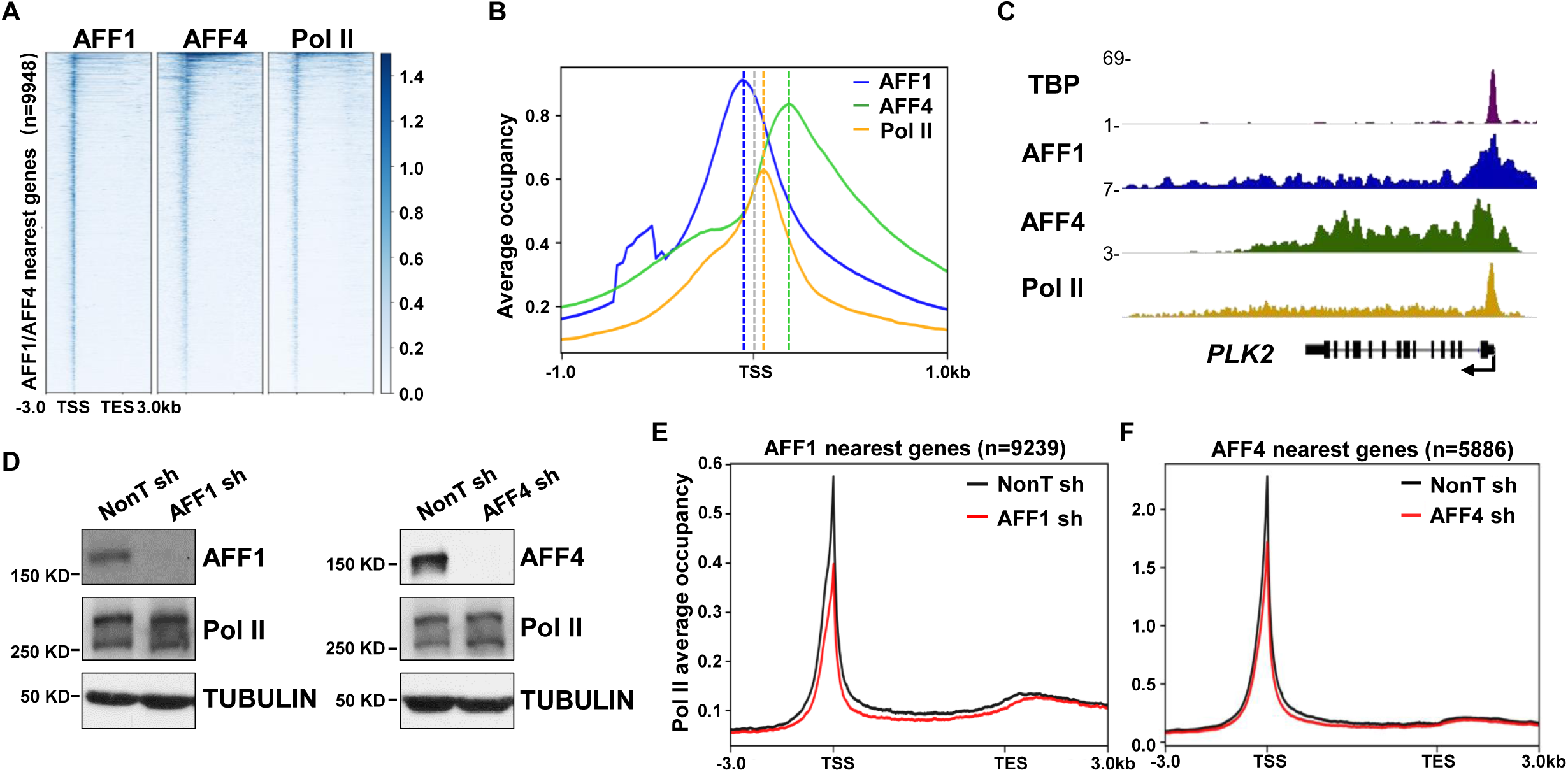
AFF1 and AFF4 exhibit diverse chromatin occupancy to regulate Pol II transcription diversely. (A) Heatmap showing AFF1, AFF4 and Pol II ChIP-seq signals at AFF1 and AFF4 nearest genes from 3 kb upstream of TSS to 3 kb downstream of transcription termination site (TES). (B) Metaplot analysis of AFF1, AFF4 and Pol II occupancy at AFF1 and AFF4 nearest genes from 1 kb upstream to 1 kb downstream of TSS. Grey horizontal dotted line indicates TSS, other three different color horizontal dotted lines indicate AFF1, AFF4 and Pol II peaks respectively. (C) Genome browser track example for the TBP, AFF1, AFF4 and Pol II occupancy profiles. (D) Western blot analyses showing the protein levels change of Pol II, AFF1 or AFF4 in AFF1 or AFF4 knockdown A549 cells. α-Tubulin was used as a loading control. (E) Metaplot of Pol II ChIP-seq signal at AFF1 nearest genes in control and AFF1 depletion cells over regions 3 kb upstream of TSS to 3 kb downstream of TES. (F) Metagene analysis comparing Pol II occupancy at AFF4 nearest genes in control and AFF4 knockdown cells from 3 kb upstream of TSS to 3 kb downstream of TES.

We next carried out Pol II ChIP-seq analysis in AFF1- and AFF4-depleted cells to investigate their potential roles in transcription regulation (Figure 1D and Figure 1— figure supplement 1C). AFF1 knockdown led to reduced Pol II levels at both promoters and transcribed genic regions, while AFF4 seemed to have minor effects on global Pol II occupancies (Figure 1E and F). Furthermore, western blot analysis revealed that total Pol II protein levels remained unchanged after AFF1 or AFF4 depletion (Figure 1D), indicating that AFF1-SEC has a direct effect on global Pol II genomic occupancies, but not through regulating Pol II protein levels. Thus, our results provided further evidence to support that AFF1 and AFF4 may function differently.

### Depletion of AFF4 leads to slow elongation and inefficient termination in a subset of active genes

We then scrutinized the effects of AFF4 on Pol II occupancies, and found a subset of genes with strong AFF4 binding signature showed a striking shift of Pol II peaks from 3’ to 5’ end around the transcriptional termination sites (TTS) upon AFF4 knockdown (Figure 2A and B). We also generated acute AFF4 degradation embryonic stem cell line using the auxin-inducible degron (AID) system and observed similar effects of AFF4 on transcriptional termination of active genes (Figure 2—figure supplement 1A-C). To further investigate the roles of AFF4 in regulating Pol II elongation, we performed precision nuclear run-on and sequencing (PRO-seq) in AFF4-depleted A549 cells to detect Pol II occupancy precisely. PRO-seq analysis confirmed that AFF4 knockdown resulted in early termination defects, which were previously observed in the slow Pol II mutants (Figure 2C). We next measured whether Pol II elongation rate was affected after AFF4 depletion. A549 cells were treated with the transcription inhibitor 5, 6-dichloro-1-β-D-ribofuranosylbenzimidazole (DRB) to inhibit elongation, then released from the elongation block by culturing in fresh DRB free medium for different time before qRT-PCR analyses. The short gene (∼6 kb) *PLK2* and the longer gene (∼29 kb) *UGDH* were used as examples to examine pre-mRNA levels at different intron-exon junctions (Figure 2D). With the DRB release time progressed, the increase of pre-mRNA levels of the examined genes, no matter short or long, slowed down obviously (Figure 2E). Thus, the depletion of AFF4 results in severe reduction in Pol II elongation rate.

**Figure 2.**
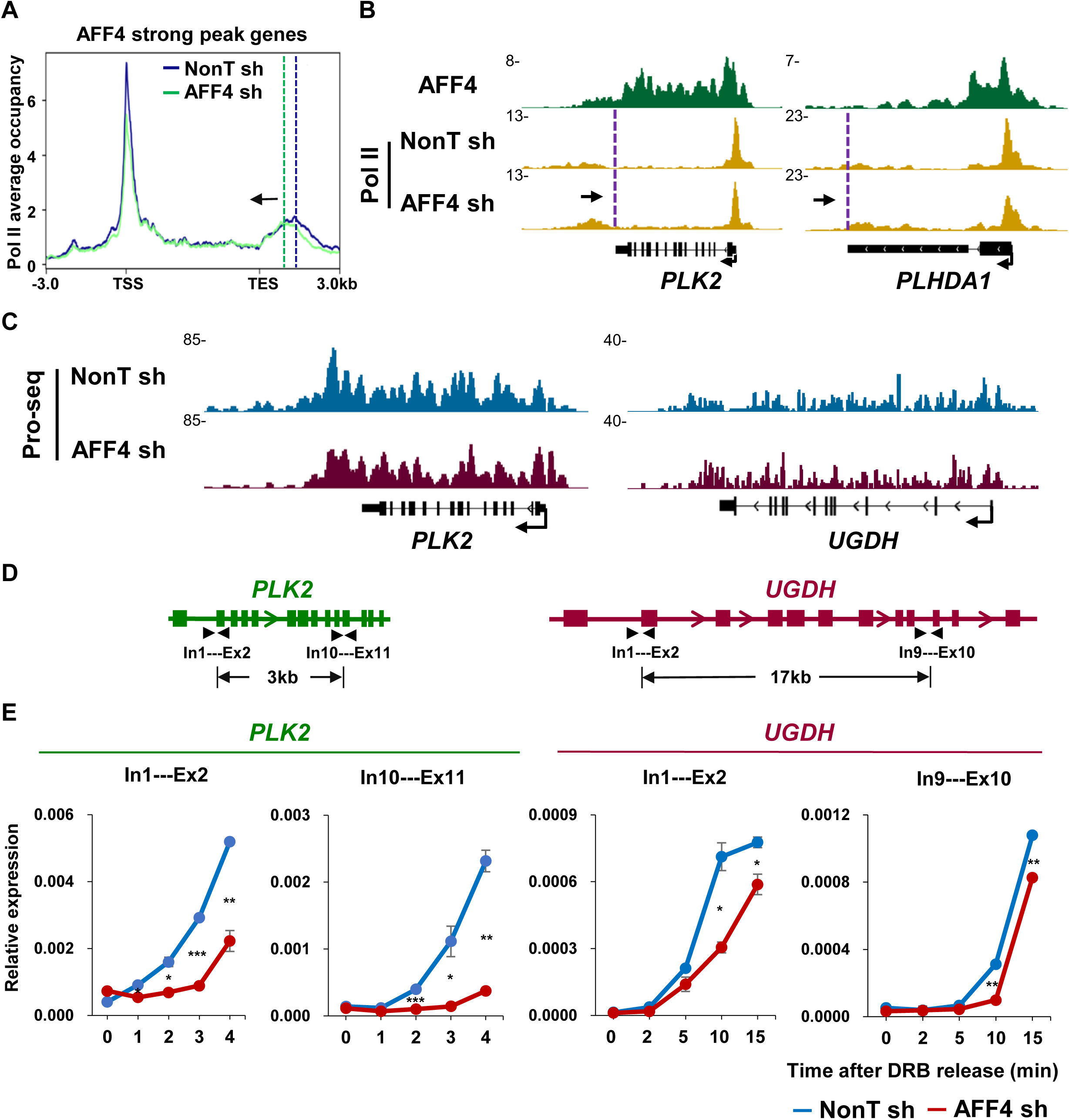
AFF4 is required for Pol II transcriptional elongation rate. (A) Metaplots of Pol II ChIP-seq read densities within −3 kb to TSS and +3 kb of TES regions in the control and AFF4 knockdown A549 cells in a subset of active genes with strong AFF4 binding peaks. Vertical dotted lines represent Pol II pause site in the termination region. Black arrows indicate the direction of Pol II peak shift. (B) Genome browser track examples of the AFF4 ChIP-seq signals, and Pol II occupancy profiles in AFF4 KD A549 cells. (C) Genome browser track examples of PRO-seq signals in wildtype and AFF4 KD A549 cells. (D) Diagrams showing the positions of pre-mRNA products at *PLK2* and *UGDH* in DRB-qRT-PCR assay. (E) qRT-PCR analysis showing pre-mRNA production in different positions of *PLK2* and *UGDH* at different DRB release times in wildtype and AFF4 knockdown A549 cells.

### AFF4 knockout leads to early termination of Pol II in the serum-inducible genes

To further examine the distinct roles of AFF1 and AFF4 in transcriptional regulation, we generated AFF1 and AFF4 KO HCT116 cells using the CRISPR-Cas9 technology (Figure 3—figure supplement 1A), and measured Pol II occupancy change after AFF1 or AFF4 knockout in the serum-inducible genes, which are well characterized for studying Pol II pause release (Lin et al., 2011). Consistently, AFF1 KO led to global reduction in Pol II levels at both promoter and genic regions, and thus impaired induction of the serum-inducible genes (Figure 3A; 3C-F; Figure 3—figure supplement 1B and D). Whereas inefficient transcriptional termination was observed in AFF4 KO cells with substantial increases in Pol II levels at transcribing units of the serum-inducible genes upon serum treatment (Figure 3B-C and Figure 3—figure supplement 1C-D). Further single molecule FISH and RT-qPCR analyses also showed a significantly increased induction of the serum-inducible genes after AFF4 KO (Figure 3D-F). Collectively, our data supported a major role of AFF1 in inducing the serum-inducible genes, while AFF4 mainly functions in transcriptional elongation step.

**Figure 3.**
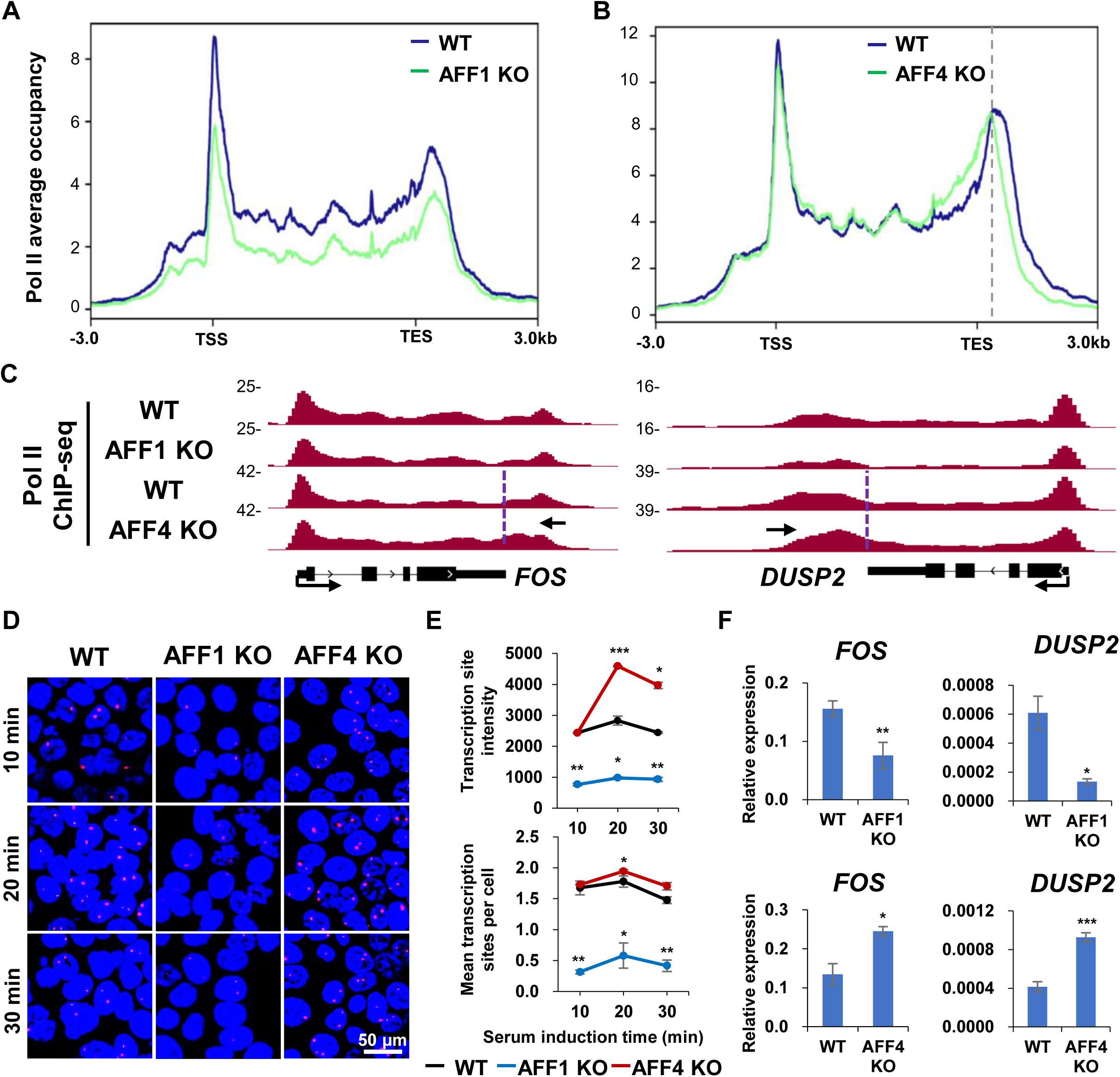
AFF4 knockout leads to early termination. (A-B) Metagene analysis comparing Pol II occupancy on serum induction genes between wildtype, AFF1 KO and AFF4 KO cell lines. Pol II density is mapping to a 6 kb window around the genes. Grey vertical dotted lines represent Pol II pause sites in termination zone. (C) Pol II ChIP-seq for *FOS* and *DUSP2* two individual genes are shown in wildtype, AFF1 KO and AFF4 KO cell lines. Purple vertical dotted lines denote the TES and black arrows indicate Pol II peak shift towards 5’ ends. (D) Images of FOS RNA FISH showing differences of nascent transcript in wildtype, AFF1 KO and AFF4 KO upon serum induction at three periods. DNA was stained with DAPI in blue. (E) Mathematical statistics analysis of mean transcription sites and transcription site intensity at *FOS* per cell after serum induction. (F) Real-time qPCR detection of *FOS* and *DUSP2* levels in WT, AFF1 KO and AFF4 KO cells.

### AFF4-P-TEFb interaction is essential for maintaining proper transcriptional elongation rate

Next, we sought to investigate the mechanism underlying Pol II inefficient termination after AFF4 depletion. The first 100aa of AFF4 is the P-TEFb interaction domain (PID) (He et al., 2011). We used the CRISPR-Cas9 technique to delete the AFF4 N-terminal region and obtained the AFF4-PID deletion (residue 18-57 deleted) cell lines (Figure 4A and Figure 4—figure supplement 1A). Immunoprecipitation assays confirmed that the AFF4-PID deletion mutant (AFF4 MT) can still interact with other SEC subunits, but not CDK9 (Figure 4B). As shown by Pol II ChIP-seq analysis, PID deletion in AFF4, similar to AFF4 KO, also led to inefficient termination (Figure 4C-D and Figure 4— figure supplement 1B-C). Single molecule FISH and RNA-seq analysis indeed showed a significantly increased induction level of the serum-inducible genes after PID deletion in AFF4 (Figure 4E-G and Figure 4—figure supplement 1D). These data indicated that the interaction of AFF4 and P-TEFb is essential for proper elongation of Pol II.

**Figure 4.**
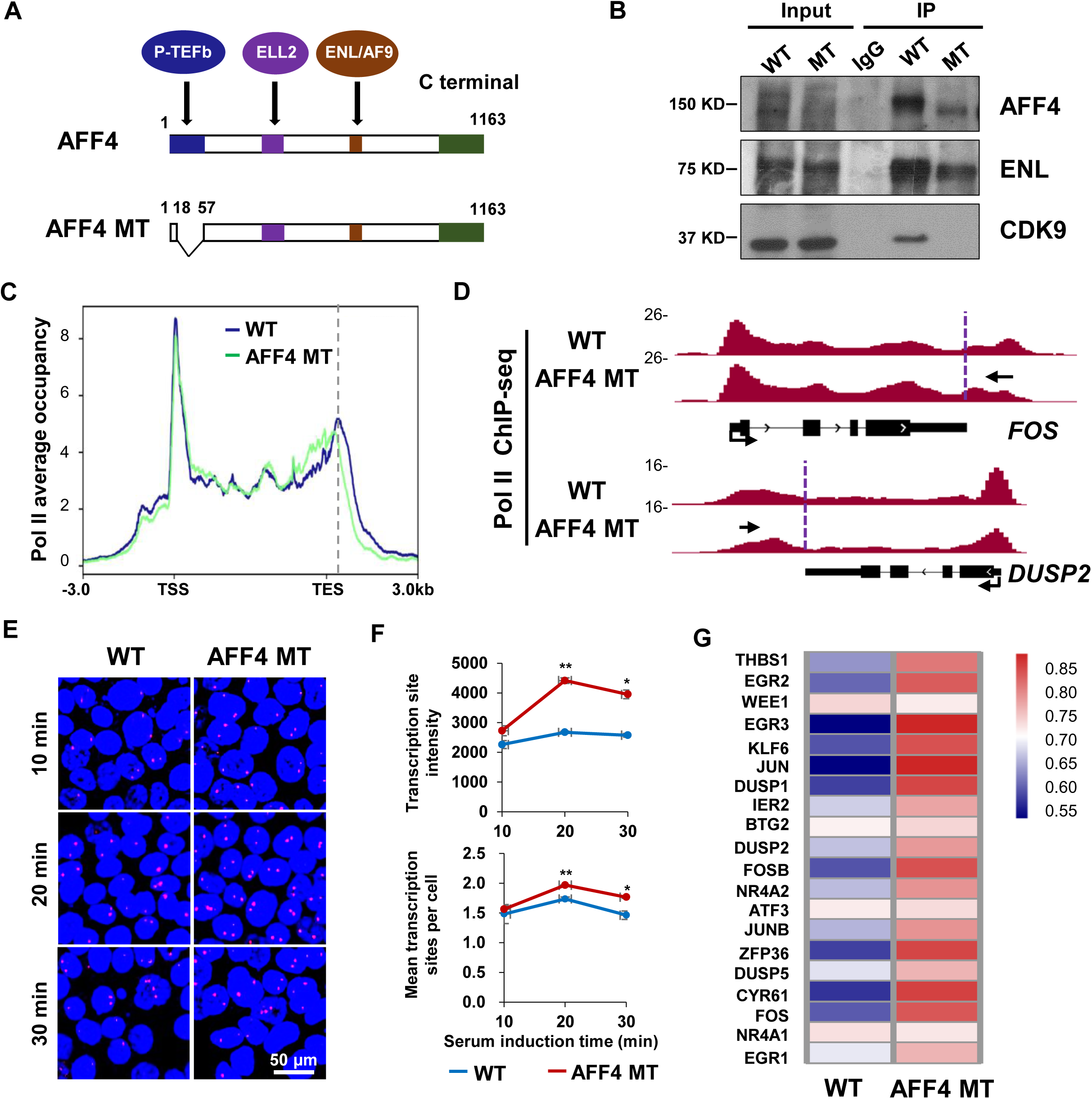
AFF4-P-TEFb interaction is important for AFF4 associated Pol II termination. (A) Schematic diagrams indicate AFF4 interaction regions and the deletion region of AFF4 mut cell. (B) Western blot analysis of AFF4, ENL and CDK9 in the wild type and AFF4 depletion as well as mutation HCT116 cells through AFF4 immunoprecipitation. (C) Metagene analysis comparing Pol II ChIP-seq signals at the serum-induced genes between wildtype and AFF4 MT cell lines. Pol II density is mapping to a 6 kb window around the genes. Grey vertical dotted lines represent Pol II pause sites in termination zone. (D) Pol II ChIP-seq for *FOS* and *DUSP2* two individual genes are shown in wildtype and AFF4 MT cell lines. Purple vertical dotted lines denote the TES and black arrows indicate Pol II peak shift towards 5’ ends. (E) Images of FOS RNA FISH showing differences of nascent transcript in wildtype and AFF4 MT cells upon serum induction at three periods. DNA was stained with DAPI in blue. (F) Mathematical statistics analysis of mean transcription sites and transcription site intensity at *FOS* per cell after serum induction. (G) Heatmap of RNA sequencing datagenerating by log_2_ fold change shows differential expression levels of serum induction genes in wildtype and AFF4 MT cells.

### AFF4 mutation results in accumulation of Ser2P, Ser5P Pol II and elongation factors in gene bodies

As both AFF4 KO and the PID deletion led to similar transcription elongation defects, we thus used the AFF4 MT lines for further study. We investigated the occupancy changes of elongation factors, including AFF1, CDK9 and SPT5, in the AFF4 MT cells. Manual ChIP-qPCR analysis showed significantly increased occupancies of AFF1, CDK9 and SPT5 at gene body, but decreased enrichment at 3’ end of the *FOS* gene in the AFF4 MT cells (Figure 5A-C). It has been reported that depletion of PAF1 or SPT6 leads to Pol II accumulation at 5’ ends of genes and early termination (Ardehali et al., 2009; Hou et al., 2019). Interestingly, we found increased occupancies of PAF1 and SPT6, together with Pol II, in the AFF4 MT cells (Figure 5D-E). It is likely that depletion of AFF4 slowed down the elongation rate of Pol II, thus leading to the accumulation of the transcriptional elongation factors.

**Figure 5.**
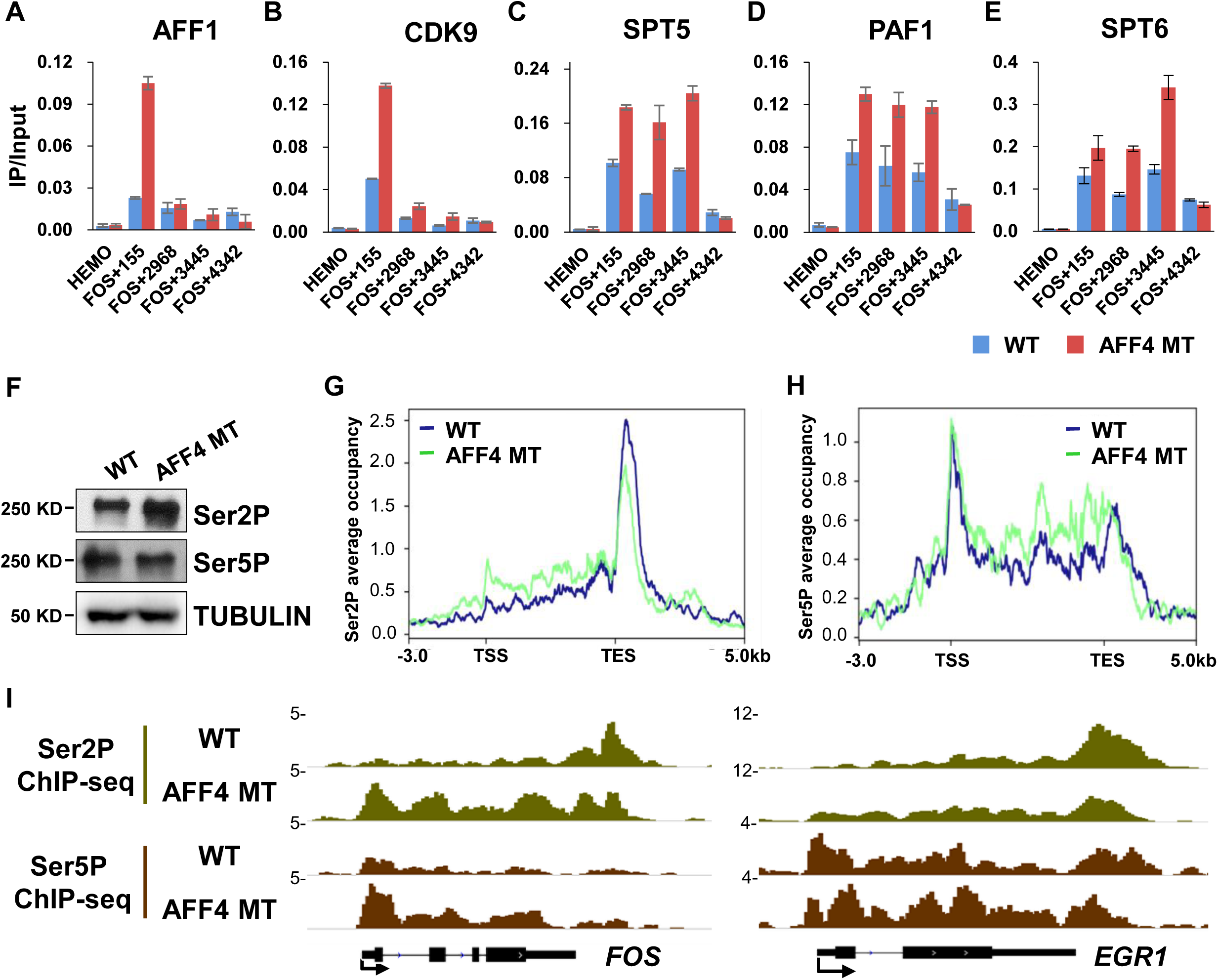
AFF4 mutation results in the accumulation of Ser2P, Ser5P Pol II and elongation associated factors at 5’ end of genes. (A-C) ChIP-qPCR analysis showed the occupancies of elongation factors (AFF1, CDK9, and SPT5) at the *FOS* gene in AFF4 mutation serum induced HCT116 cells. (D-E) ChIP-qPCR analysis showing that the elongation factors PAF1 and SPT6 occupancy change at the *FOS* gene in control and AFF4 mutant cells. (F) The protein levels of Ser2P and Ser5P in AFF4 mutation HCT116 cells. α-Tubulin was used as a loading control. (G-H) Metaplot analysis of Ser2P and Ser5P ChIP-seq for the serum induced genes in AFF4 MT cell line. Ser2P and Ser5P density are plotted in a −3 kb and +5 kb window around the genes. (I) Genome browser tracks of Ser2P and Ser5P ChIP-seq at induction genes *FOS* and *EGR1* in control and AFF4 mutation cells in HCT116.

Previous studies have reported that Ser2P Pol II is accumulated at 5’ end of genes in the slow Pol II mutant (Fong et al., 2017). AFF4 depletion leads to early termination, similar to the phenotype of the slow Pol II mutant (Fong et al., 2015). Notably, we observed that the protein levels of Ser2P Pol II were increased dramatically in the AFF4 MT cells (Figure 5F and Figure 5—figure supplement 1A). Consistently, Ser2P Pol II increased around TSS and gene bodies while significantly decreased at termination sites in the AFF4 MT cells (Figure 5G; 5I and Figure 5—figure supplement 1B). Likewise, accumulation of Ser5P Pol II in the 5’ region of genes was also observed (Figure 5H-I and Figure 5—figure supplement 1C). Taken together, AFF4-P-TEFb interaction is essential for the proper Pol II elongation, but has no obvious effect on the recruitment of other SEC components and key elongation factors PAF1 and SPT6.

### AFF4 disruption leads to increased CSTF2 occupancy with a 5’ shift

In eukaryotes, CPSF, CstF and the cleavage complexes are mainly responsible for the transcription termination of protein-coding genes (Eaton and West, 2020). We further examined the protein levels of pre-mRNA 3’ end processing factors and termination factors in AFF4 KO cells and observed an obvious increase of the CstF components CSTF2 protein levels in AFF4 KO and PID deletion cells (Figure 6A and Figure 6—figure supplement 1A). However, other tested factors, such as CPSF3, SYMPK, XRN2, PPP1CB, PPP1CC, remained unchanged in AFF4 KO and PID deletion cells (Figure 6A and Figure 6—figure supplement 1A). We then examined the occupancy of CSTF2 at transcription termination regions when AFF4 is depleted (Figure 6B). Manual ChIP-qPCR analysis indicated increased enrichment with 5’ shift of CSTF2 at the TTS of the *FOS* gene in AFF4 MT cells (Figure 6B). To further examine whether the increased CSTF2 levels could contribute to the early termination defects in AFF4 mutants, CSTF2 was knockdowned in these cells (Figure 6C). Interestingly, upregulation of FOS mRNA and increase in Pol II occupancies at the *FOS* gene in AFF4 mutant were significantly compromised upon CSTF2 depletion (Figure 6D-E). Similar effects were also observed after CPSF3 depletion (Figure 6—figure supplement 1B-D). Taken together, our data demonstrated an important role of AFF4 in maintaining proper transcriptional elongation and termination.

**Figure 6.**
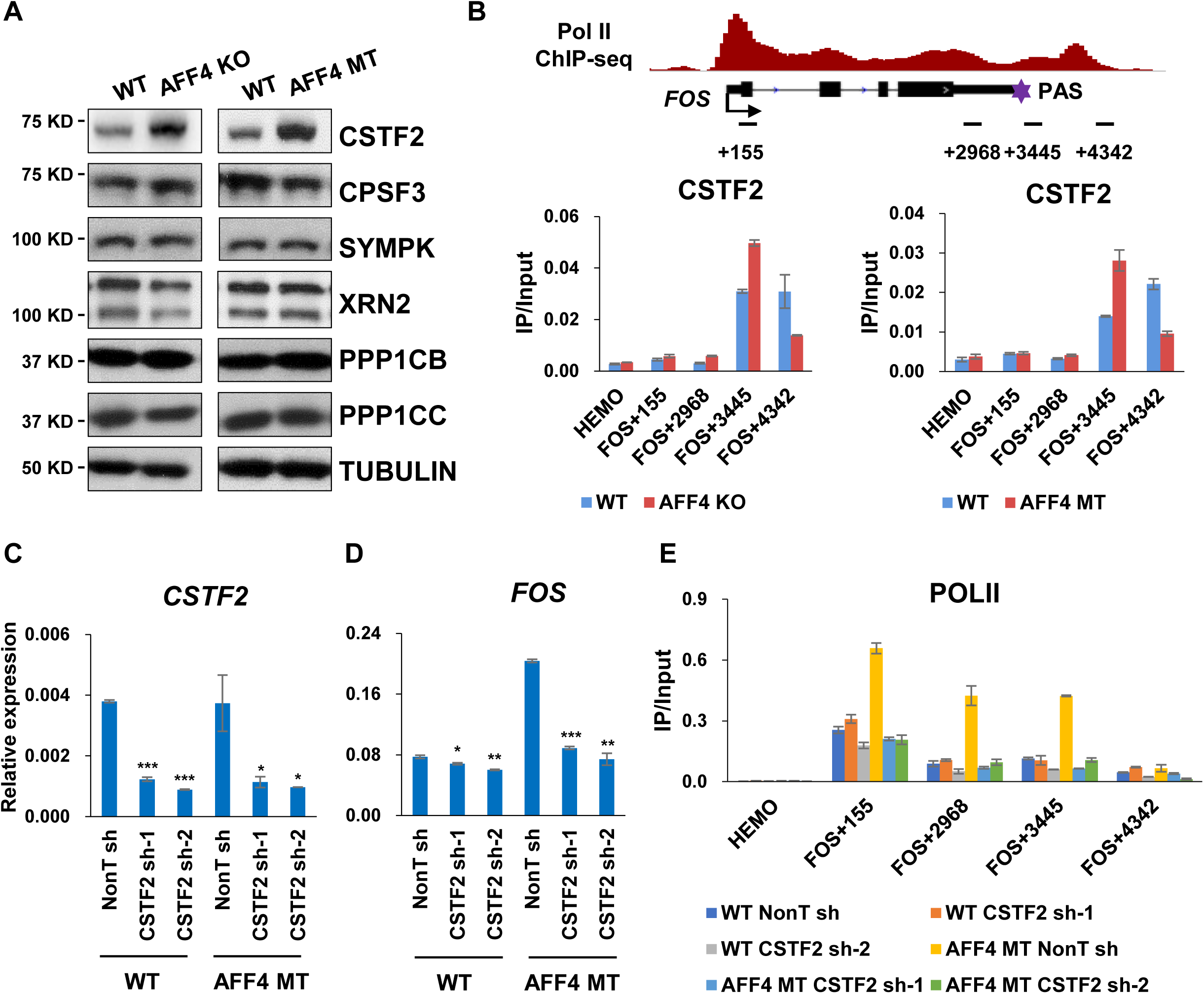
AFF4 disruption leads to increased CSTF2 occupancy with a 5’ shift. (A) The protein levels of CPF components CSTF2, CPSF3, SYMPK, Xrn2, PPP1CB and PPP1CC in AFF4 depletion and AFF4 mutation HCT116 cells. α-Tubulin was used as a loading control. (B) Schematic diagram shows the primers sites around *FOS* gene. ChIP-qPCR analysis showing the occupancy of CSTF2 at *FOS* gene in AFF4 depletion and AFF4 mutation serum induced HCT116 cells. The *HEMO* gene acts as a negative control for ChIP-qPCR. (C) RT-qPCR analysis of *CSTF2* gene expression after CSTF2 knockdown in wildtype and AFF4 mutation HCT116 cell lines. (D) RT-qPCR analysis of *FOS* gene expression after CSTF2 knockdown in wildtype and AFF4 mutation HCT116 cell lines. (E) ChIP-qPCR showing the occupancy of Pol II after CSTF2 knockdown in AFF4 mut. The *HEMO* gene acts as a negative control for ChIP-qPCR.

### AFF4 occupancies are substantially increased at a subset of highly active genes after AFF1 depletion

We noticed that there was a significant increase of AFF4 protein levels after AFF1 knockdown, and *vice versa* (Figure 7A and Figure 7—figure supplement 1A). Interestingly, we also found that AFF1 knockdown led to readthrough of Pol II around the TTS (Figure 7B-C). The Pol II defect in AFF1 depleted cells was reminiscent of fast Pol II phenotypes (Fong et al., 2015). RT-qPCR analyses after release from DRB treatment further revealed that with the DRB release time progressed, UGDH pre-mRNA increased earlier after AFF1 depletion at the Intron1-Exon2 and Intron9-Exon10 junctions, while there were no obvious changes at the short gene *PLK2* (Figure 7D). We thus speculated that the elongation rate and readthrough defects in AFF1-depleted cells could possibly result from the increased levels of AFF4. Indeed, ChIP-seq analysis confirmed the significant increased AFF4 levels at these genes, including the *PLK2* and *PHLDA1* genes (Figure 7E-F). Taken together, these data could point out a major role of AFF4-SEC in controlling the elongation rate of Pol II.

**Figure 7.**
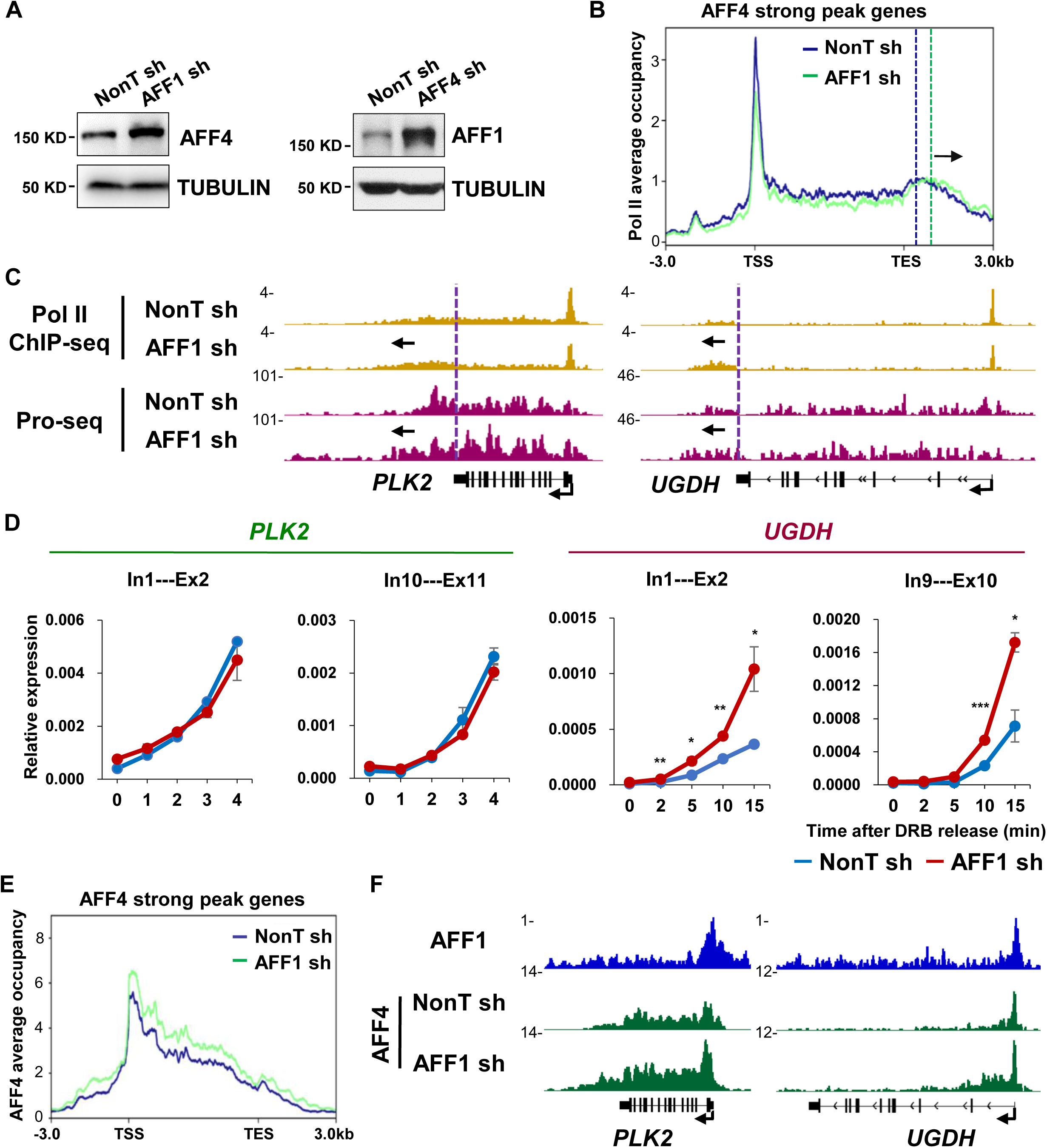
Significant increases of AFF4 occupancies at a subset of AFF1-depleted highly active genes. (A) Western blot analyses showing the protein level change of AFF4 in AFF1 knockdown and AFF1 in AFF4 knockdown A549 cells. α-Tubulin was used as a loading control. (B) Metaplots of Pol II ChIP-seq read densities within −3 kb to TSS and +3 kb of TES regions in the control and AFF1 knockdown A549 cells in the subset of active genes with strong AFF4 binding peaks. Vertical dotted lines represent Pol II pause site in the termination region. Black arrows indicate the direction of Pol II peak shift. (C) Genome browser track examples of the Pol II ChIP-seq signals and pro-seq signals in wildtype and AFF1 KD A549 cells. Purple vertical dotted lines denote the TES and black arrows indicate Pol II peak shift towards 3’ ends. (D) qRT-PCR analysis showing pre-mRNA production in different positions of *PLK2* and *UGDH* at different DRB release times in wildtype and AFF1 knockdown A549 cells. (E) Metaplots showing that the increased AFF4 occupancy in the subset of active genes with strong AFF4 binding peaks. The analysis area is −3 kb of TSS and +3 kb of TES. (F) Genome browser track examples of the AFF1 ChIP-seq, and AFF4 occupancy profiles in control and AFF1 KD cells.

## Discussion

Promoter-proximal pausing of RNA Pol II is a prevalent and critical step in eukaryotic transcription (Zhou et al., 2012; Chen et al., 2018; Core and Adelman, 2019; Schier and Taatjes, 2020). The activity of P-TEFb is tightly controlled *in vivo* through forming different complexes, indicating the regulatory complexity of paused Pol II (Lu et al., 2016; Li et al., 2018; Bacon and D’Orso, 2019). AFF1 and AFF4 share similar domain structure and serve as scaffolds for SEC assembly (Luo et al., 2012a). Yet, it is still unclear regarding their specific roles in Pol II transcriptional elongation control. Here, we revealed the different genome occupancy profiles with AFF1 and AFF4 peaking at upstream and downstream of TSS, respectively. AFF1 is more inclined to function at the transcriptional initiation step and depletion of AFF1 significantly reduces Pol II levels at both promoter and transcribing unit. AFF4 tends to serve as an elongation rate monitor to ensure the proper transcription elongation (Figure. 8).

**Figure 8.**
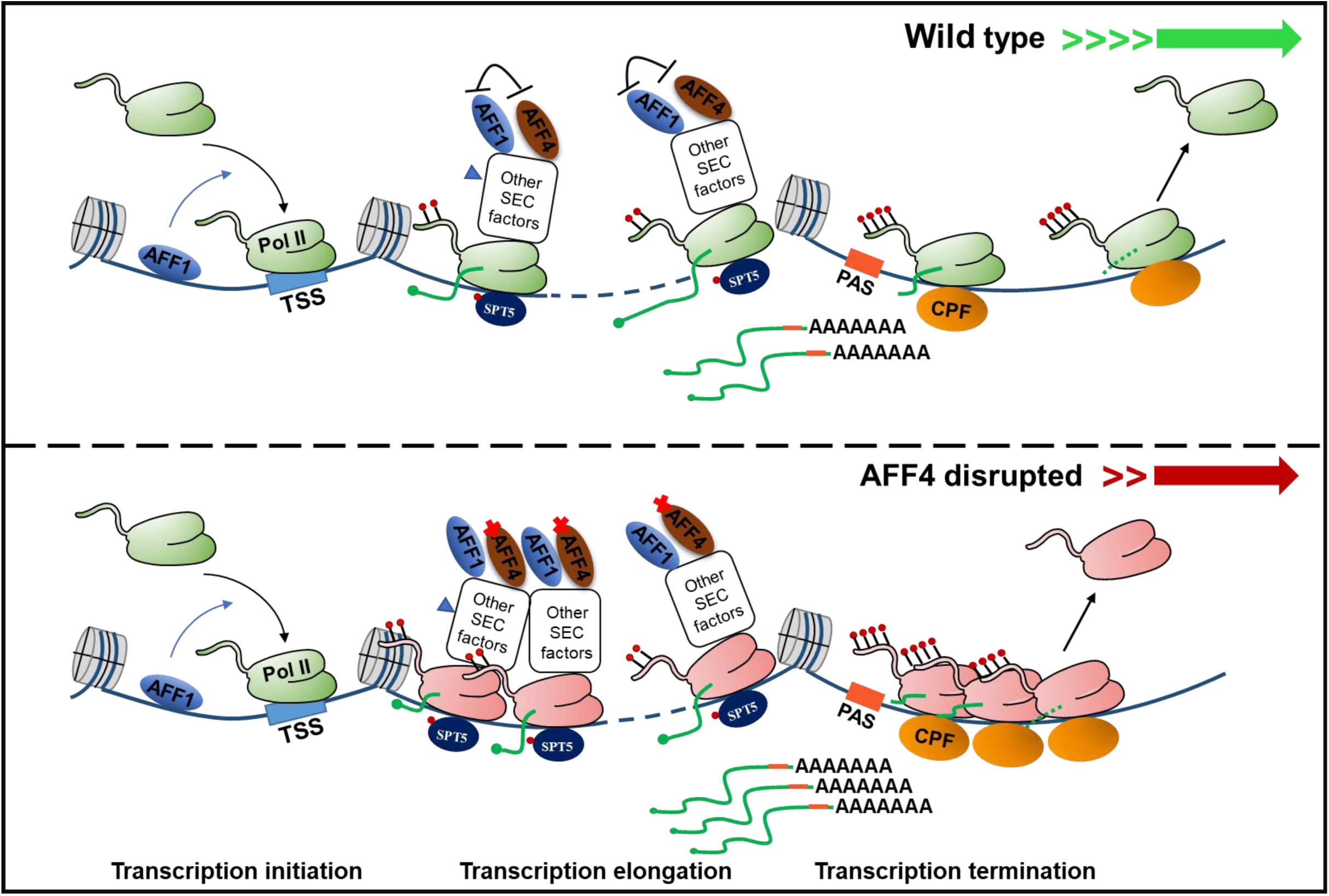
Cartoon model indicates the roles of AFF1 and AFF4 in regulating Pol II transcription. AFF1 binds upstream of TSS, while AFF4 is enriched downstream of TSS. In a subset of active genes with strong AFF4 binding signature, AFF4 disruption causes slow elongation and early termination.

Substantial progresses have been made in understanding the functions of most of the individual SEC subunits. AFF1 and AFF4 share similar domain structure and serve as scaffolds for SEC assembly (Lin et al., 2010). SEC inhibition by the small molecular inhibitors KL-1/2, which destabilize both AFF1 and AFF4, blocks the release of paused Pol II at the genome-wide scale (Liang et al., 2018). It was proposed that AFF1 and AFF4 are able to form either heterodimer or homodimer via their C-terminal homology domains (CHDs), thus regulating different subsets of genes (Yokoyama et al., 2010; Lu et al., 2015). We and others have previously shown that P-TEFb within AFF4-SEC, but not AFF1-SEC, is essential in promoting the release of Pol II from pausing to active elongation at the heat shock responsive gene *HSP70* and the *c-MYC* gene (Lin et al., 2010; Takahashi et al., 2011; Luo et al., 2012a). Our data here demonstrated that AFF1 and AFF4 co-occupy the highly transcribed genes in A549 cells. However, only AFF1, but not AFF4, within SEC plays a major role in promoting pause release at the highly transcribed genes. The reduction of Pol II levels was observed not only at gene bodies, but also at promoter regions after depletion of AFF1. A recent study reported that the pSER domain of AFF1 is able to facilitate Pol II mediated transcriptional initiation through interacting with the TATA-binding protein (TBP) loading factor selectivity factor 1 (SL1) (Okuda et al., 2015). It is likely that AFF1 could stimulate transcription at initiation and early elongation stages via different protein complexes at the highly transcribed genes.

Interestingly, AFF4 enriches at ∼175nt downstream of TSS and depletion of AFF4 results in slow elongation and early termination in a subset of active genes and especially at the serum-inducible genes, mirroring the effects of the slow Pol II mutant. Our study indicated that unlike AFF1, AFF4 could mainly function at the elongation step. SEC mediates rapid transcriptional induction of the serum-inducible genes. Previous studies have shown that the depletion of some elongation factors, such as PAF1 and SPT6, result in slow elongation rate (Ardehali et al., 2009; Hou et al., 2019). However, we failed to observe the decreased occupancy of these factors when AFF4 disrupted, suggesting that AFF4 regulates elongation rate in other ways. In AFF4 KO cells, we also observed a clear 5’ switch of Pol II Ser2P and termination factors, like CSTF2, at the serum-inducible genes upon serum treatment. Similar phenotype was observed in the AFF4 MT cell line, indicating that AFF4 regulates transcription depending on its interaction with P-TEFb to a large extent. It needs to be further examined the potential mechanism underlying early termination defects after AFF4 depletion.

## Materials and Methods

### Cell lines and culture conditions

A549, HCT116, HEK293T and AFF1 KO/AFF4 KO/MT HCT116 stable cell lines were cultured in DMEM (HyClone) supplemented with 10% FBS (Ex-Cell Bio) and 1% Penicillin-Streptomycin (HyClone), at 37°C with 5% CO_2_ in a humidified incubator and routinely passaged 2-3 times a week. Auxin inducible degron ES cell line was cultured in N2B27 medium supplemented with 2i/LIF in 0.1% gelatin-coated tissue culture flask (Ying et al., 2008).

### Plasmid construction

Human AFF1 and AFF4 sgRNA oligos were subcloned into lentiCRISPR v2 vector using Bbs1 sites. Human AFF1, AFF4, CSTF2, CPSF3 shRNA oligos were subcloned into pLKO.1 vector using Age1/EcoR1 sites. Mouse AFF4 sgRNA loigos was subcloned into px330 vector using Bbs1 site. Mouse AFF4 homology arms was cloned at BamH1 site into AID-mClover-Neo vector.

### Generation of stable cell lines

To generate AFF1/AFF4 knockout cell lines, HEK293T cells were co-transfected with two sgRNA plasmids, psPAX2 packaging plasmids and pMD2.G envelope plasmids using Lipofectamine 2000 (Thermo Fisher Scientific) according to manufacturer’s protocol. After 16 hours of transfection, the medium was replaced with the fresh medium. The lentiviral supernatants were collected 48 and 72 hours after the transfection, filtered through 0.45 μm filters. HCT116 cells were infected with filtered lentiviral supernatants together with polybrene (Sigma) at the concentration of 8 μg/ml. After 24 hours, the infected cells were subjected to selection with 2 μg/ml of puromycin for 7 days. The selective medium was exchanged every 4 days. After that, colonies were washed with PBS, picked using yellow tips and transferred to 48-well plate containing selective medium. When the cells grew full, a half of the cells were used for extracting genomic DNA and the reminding were frozen. To generate AFF4-AID ES cell line, CRISPR/Cas and donor plasmids were transfected in six-well OSTIR1-expressing ES cell line using HighGene (Abclonal). Two days after trasfection, the cells were selected in the presence of 300 μg/ml G418 (Sigma). After a week, the single colony was picked to grow to identify (Natsume et al., 2016).

### Immunoprecipitations and western blot

Cells were harvested in cold PBS and pelleted by centrifugation at 1000 rpm for 5 minutes at 4 ℃. Then cells were lysed in high salt buffer containing 20 mM HEPES [pH 7.4], 1 mM MgCl_2_, 10 mM KCl and 420 mM NaCl adding protease inhibitor cocktail (Sigma-Aldrich) for 30 minutes at 4 ℃. The lysate was centrifugated at 14800 rpm for 15 minutes at 4 ℃. The supernatant was diluted with cold PBS to reduce the concentration of NaCl to 200 mM. Then the diluted supernatant was incubated with protein A beads and antibody for 5 hours at 4 ℃. After the incubation, the beads were washed for three times using wash buffer. Eluted proteins by boiling beads with SDS were subjected to SDS-PAGE gels and transferred to polyvinylidene fluoride membrane for western blot.

### RT-qPCR and RNA-seq library preparation

Total RNA was extracted by using the RNeasy kit (Qiagen) according to the instructions of the manufacturer. RNA was treated with DNase I (NEB) and re-purified on the column. Reverse transcription was performed by PrimeScript™ RT Master Mix (TaKaRa). cDNA was amplified using iTaq™ Universal SYBR® Green Supermix (Bio-Rad) on CFX96 (Bio-Rad). Housekeeping gene *GAPDH* were used as normalization control. To generate RNA-seq library, total RNA was polyadenylated, fragmented and reversed. Then reversed cDNA was ligated to adapters for sequencing (Illumina).

### ChIP and ChIP-seq library preparation

ChIP was performed according to the previously published protocol (Guo et al., 2020). Briefly, A549 and HCT116 cells were crosslinked with 1% formaldehyde for 10 minutes and quenched with glycine for 5 minutes at the room temperature. Fixed cells were sonicated and immunoprecipitated with the mixture of beads and antibody overnight at 4 ℃. ChIP-DNA was purified using PCR purification kit (Qiagen) and performed for qPCR using iTaq™ Universal SYBR Green Supermix (Bio-Rad) on CFX96 (Bio-Rad). The ChIP-seq sample preparation kit was used to prepare generate ChIP-seq library.

### Antibodies

AFF1, AFF4, CDK9, ENL and Pol II antibodies were homemade antibodies (Lin et al., 2010; Yue et al., 2021). Ser2P Pol II (ab5095), Ser5P Pol II (ab5131), PAF1 (ab20662) rabbit polyclonal antibody were purchased from Abcam; SPT5 (A9193), SPT6 (A16434), CSTF2 (A8116), CPSF3 (A12368), SYMPK (A8722), Xrn2 (A18350), PPP1CB (A13528), PPP1CC (A4025) rabbit polyclonal antibody were purchased from ABclonal.

### RNA FISH and data analysis

FOS RNA FISH was performed according to the previously published protocol (Guo et al., 2020). Briefly, cells were seed on 12-mm round coverslips in a 12-well plate, after serum induction, cells were fixed by 3.7% formaldehyde and permeabilized by 70% ethanol. After washing by wash buffer A, the cells were hybridized with FISH probes in hybridization buffer for 4 hours at 37℃. Then the coverslips were mounted by mounting buffer and images were pictured by Zeiss LSM 700 confocal microscope.

### Precision Nuclear Run-on, sequencing and analysis

Precision Nuclear Run-on and Sequencing was performed according to the published protocol (Mahat et al., 2016). About 10^7^ cells were harvested by scraping and centrifuging, then, the cells were washed by cold PBS. The cells were resuspended in 10 ml cold douncing buffer, incubated for 10 minutes on ice and dounced 25 times. 2 biotinylated nucleotides were used to perform the nuclear run-on reaction for 3 minutes at 37℃. TRIzol LS (Invitrogen) was used to stop the reaction, then, the biotinylated RNA was extracted, fragmented, adapters ligated and enrichened for library construction. The libraries were size selected by 2.2% agarose gel from 140-350 bp. Clean reads were aligned to the Human genome (UCSC genome, GRCh37/hg19) using bowtie (v1.3.0) with parameter ‘-m 1 -v 2 −1’. BigWig files were created by DeepTools.

### DRB-RT-qPCR

Cells were treated with 100 μM 5, 6-dichloro-1-β-D-ribofuranosylbenzimidazole (DRB) to inhibit Pol II transcription elongation. After the incubation for 3 hours at 37℃, cells were washed with PBS three times. Then, the cells were incubated with warm medium for different time points. Cells were lysed in TRIzol reagent (Vazyme), reverse transcription was performed by ABScript III RT Master Mix (ABclonal).

### RNA-seq analysis

The clean reads were subjected to TopHat and Cufflinks pipeline based on Human genome (UCSC genome, GRCh37/hg19) (Trapnell et al., 2009; Trapnell et al., 2010; Trapnell et al., 2012). The Cuffdiff program within Cufflinks was used to test statistically significant differences of genes expressions between control and AFF4 mutant cells.

### ChIP-seq analysis

TBP ChIP-seq data in HepG2 cells was downloaded from GSE31477. AFF1, AFF4 and Pol II ChIP-seq data in A549 cells, Pol II, Ser2P Pol II and Ser5P Pol II ChIP-seq data in serum-treated HCT116 cells were generated in this study, and acquired through the default Illumina pipeline using Casava v1.8. Clean reads were aligned to the Human genome (UCSC genome, GRCh37/hg19) using Bowtie2 (v2.2.5) with default mode (Langmead and Salzberg, 2012). Peak calling was performed with MACS2 (v2.1.1) (Zhang et al., 2008). For AFF1 and AFF4 peaks, associated control samples were used to determine statistical enrichment at a p < 1e-8 and FDR < 0.05. For Pol II peaks, the enrichments were determined at p < 1e-8 and FDR < 0.05. The ChIP-seq enrichment is displayed as reads per million (RPM). Gene annotations and transcript start site information were from Human genome (UCSC genome, GRCh37/hg19) by R package ChIPseeker (Yu et al., 2015). Gene ontology (GO) analysis was performed with DAVID (Huang da et al., 2009).

### Data availability

ChIP-seq and RNA-seq data in this study have been deposited at GEO under the accession number GSE207320. TBP ChIP-seq is from GEO accession number GSE31477.

## Acknowledgments

The authors are grateful to the Lin & Luo lab members for helpful discussion of this study. We are grateful to Dr. Xiong Ji for OSTIR1-expressing ES cell line.

## Supplementary Figure Legends

**Figure 1—figure supplement 1.**
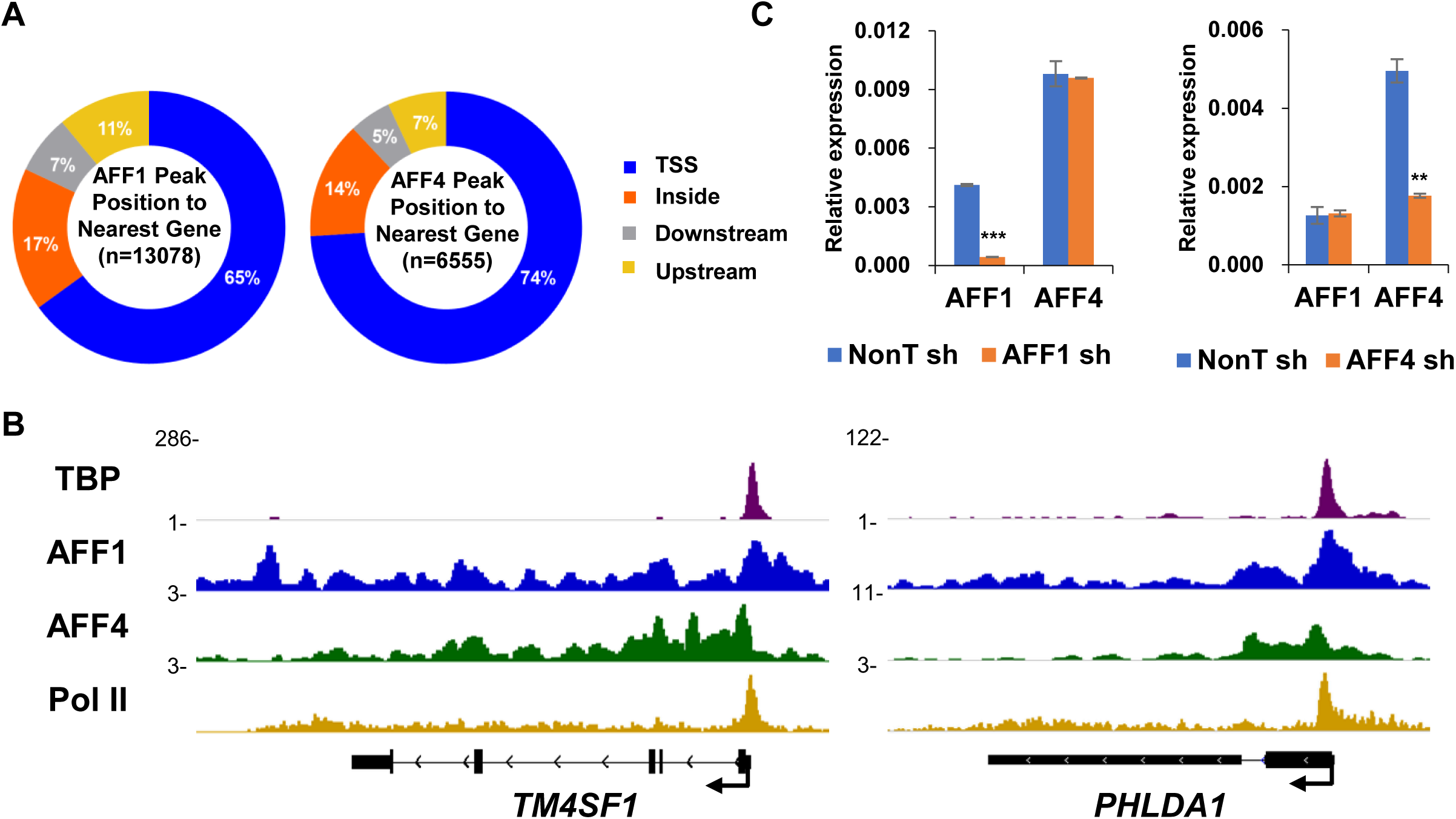
AFF1 and AFF4 exhibit diverse chromatin occupancy to regulate Pol II transcription diversely. (A) Pie charts showing that the distribution percentages of AFF1 or AFF4 peaks at different genome locations including TSS, within a gene, and upstream or downstream of the nearest gene. (B) Genome browser track examples for the TBP, AFF1, AFF4 and Pol II ChIP-seq signals. (C) RT-qPCR analyses showing the *AFF1* and *AFF4* knockdown efficiency as well as the mRNA levels of AFF1 and AFF4 were not affected by each other. Expression levels were normalized to *Gapdh*.

**Figure 2—figure supplement 1.**
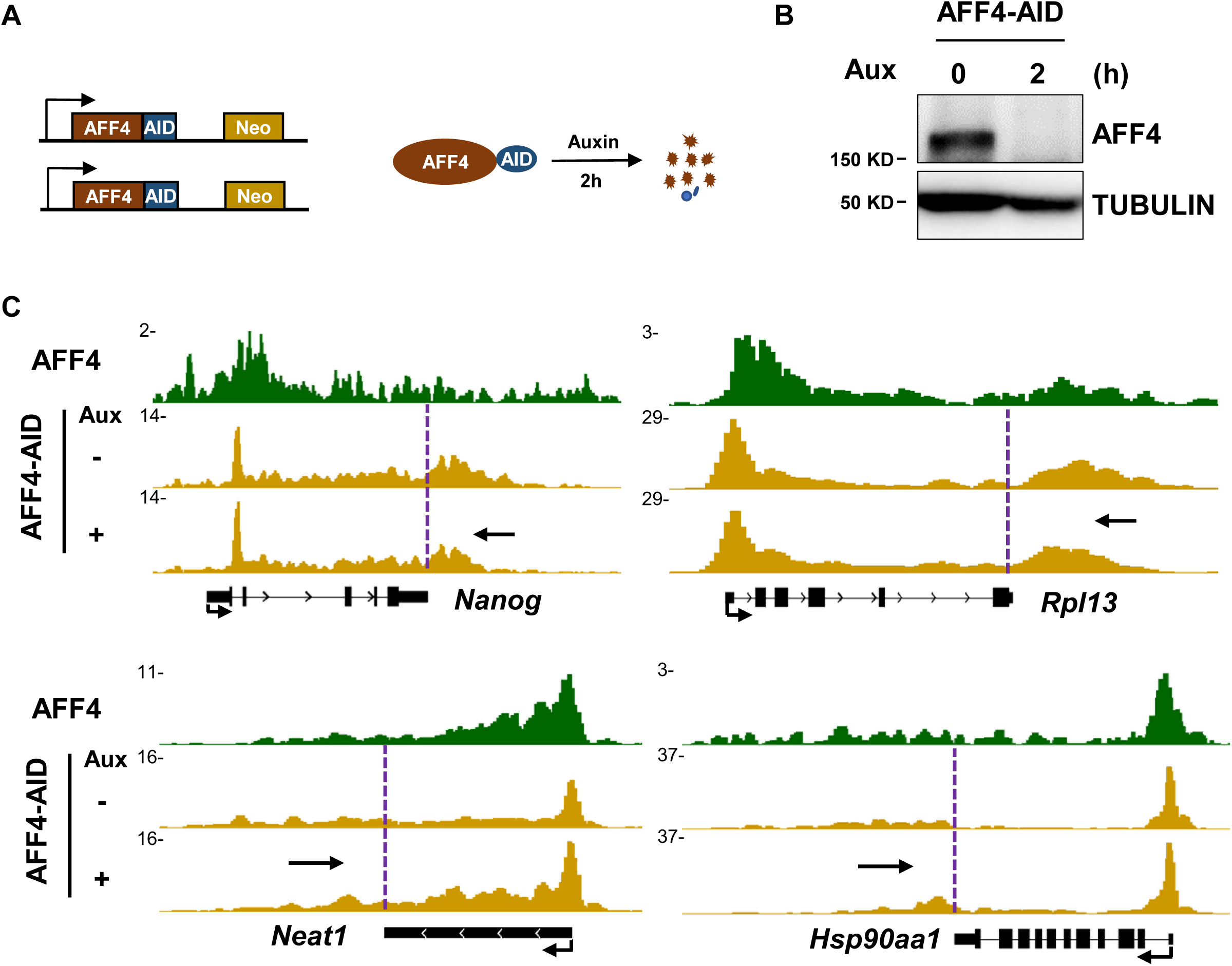
AFF4 is required for Pol II transcriptional elongation rate. (A) Schematic diagrams showing the generation and the degradation of AFF4 degron ES cells. (B) Western blot detecting the AFF4 protein levels in the AFF4 degron ES cells. The degron cells were treated without or with 500 μM Auxin for 2h. α-Tubulin was used as a loading control. (C) Genome browser track examples of the AFF4 ChIP-seq signals in embryonic stem cells, and Pol II occupancy profiles at active genes before and after auxin treatment in AFF4 degron ES cells for 2h.

**Figure 3—figure supplement 1.**
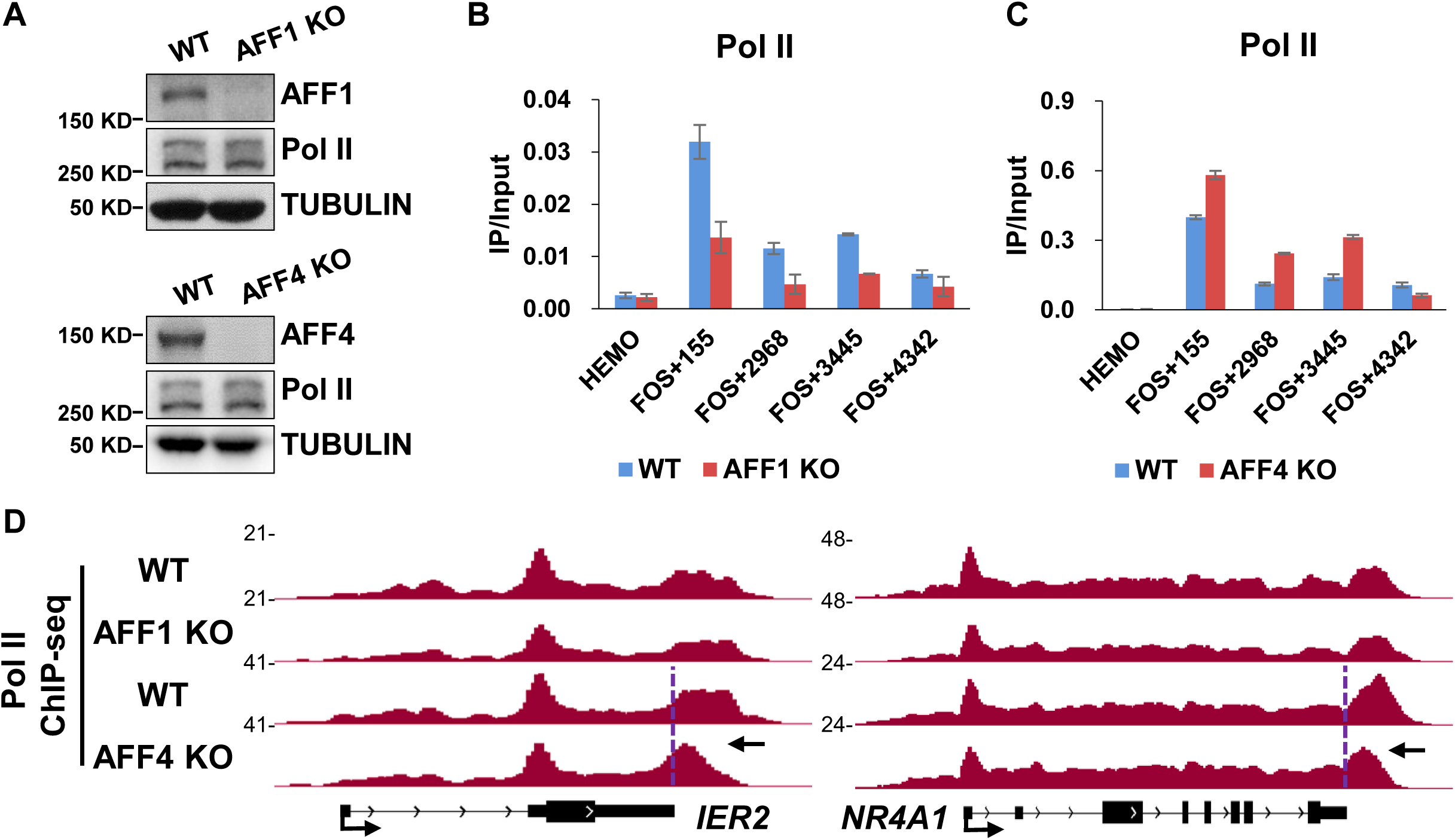
AFF4 knockout leads to early termination. (A) Western blot analyses showing the protein levels of AFF1 and Pol II in AFF1 knockout HCT116 cells as well as the protein levels of AFF4 and Pol II in AFF4 knockout HCT116 cells. α-Tubulin was used as a loading control. (B) ChIP-qPCR analysis showing that the Pol II occupancy change at *FOS* gene in control and AFF1 KO cells. (C) ChIP-qPCR analysis showing that the Pol II occupancy change at *FOS* gene in control and AFF4 KO cells. (D) Genome browser track examples of Pol II occupancy in control, AFF1 KO and AFF4 KO cells. Purple vertical dotted lines denote the TES and black arrows indicate Pol II peak shift towards 5’ ends.

**Figure 4—figure supplement 1.**
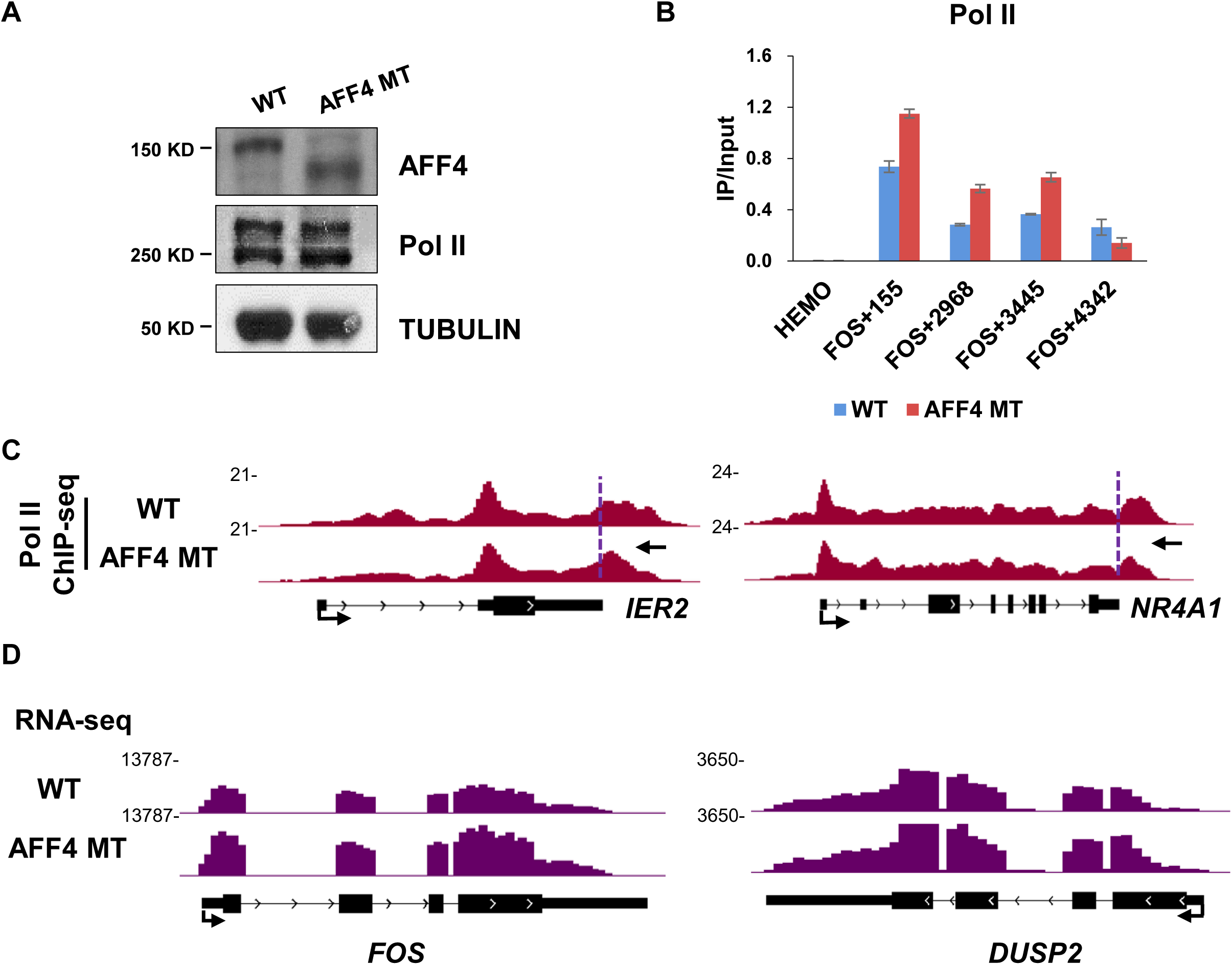
AFF4-P-TEFb interaction is important for AFF4 associated Pol II termination. (A) Western blot analyses showing the protein levels of AFF4 and Pol II in AFF4 mutation HCT116 cells. α-Tubulin was used as a loading control. (B) ChIP-qPCR analysis showing that the Pol II occupancy change at *FOS* gene in control and AFF4 MT cells. (C) Genome browser track examples of Pol II occupancy in control and AFF4 MT cells. Purple vertical dotted lines denote the TES and black arrows indicate Pol II peak shift towards 5’ ends. (D) Two individual gene tracks of RNA-seq in WT and AFF4 mut are shown.

**Figure 5—figure supplement 1.**
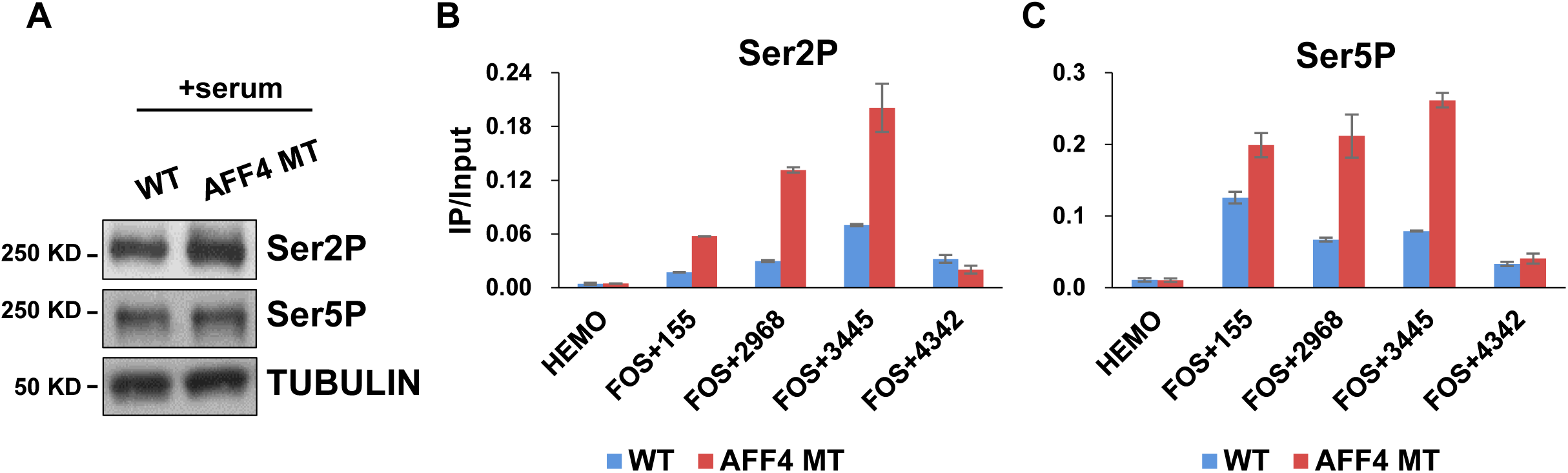
AFF4 mutation results in the accumulation of Ser2P, Ser5P and elongation associated factors at 5’ end of genes. (A) The protein levels of Ser2P and Ser5P Pol II in AFF4 mutation HCT116 cells under serum induction. α-Tubulin was used as a loading control. (B) ChIP-qPCR analysis showing that the Ser2P Pol II occupancy change at the *FOS* gene in control and AFF4 mut serum induced HCT116 cells. (C) ChIP-qPCR analysis showing that the Ser5P occupancy change at the *FOS* gene in wildtype and AFF4 MT serum induced HCT116 cells.

**Figure 6—figure supplement 1.**
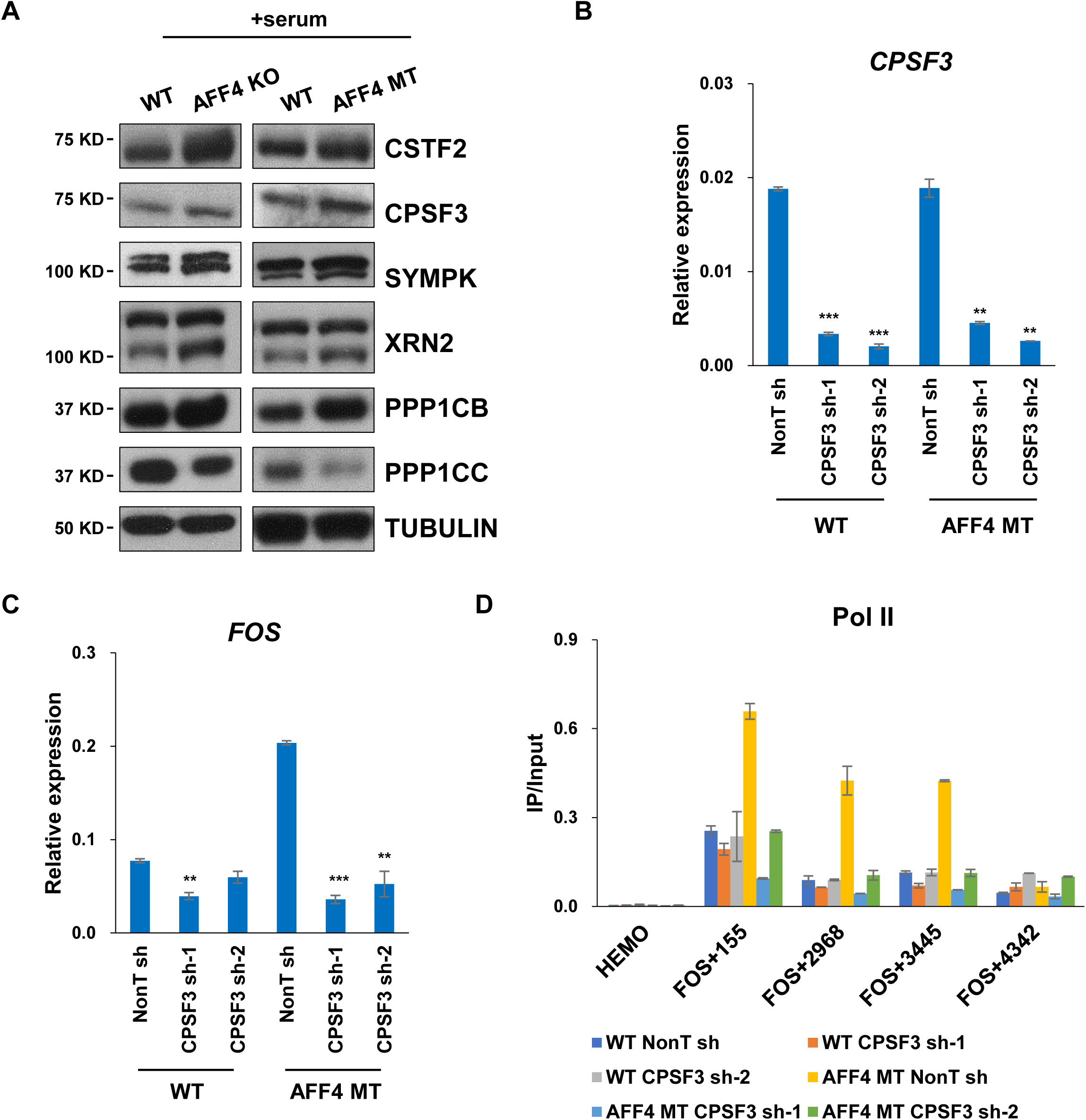
AFF4 disruption leads to increased CSTF2 occupancy with a 5’ shift. (A) The protein levels of CPF components CSTF2, CPSF3, SYMPK, Xrn2, PPP1CB and PPP1CC in AFF4 depletion and AFF4 mutation serum induced HCT116 cells. α-Tubulin was used as a loading control. (B) RT-qPCR showing the RNA expression level of *CPSF3* after CPSF3 knockdown in wildtype and AFF4 MT cells. (C) RT-qPCR showing the RNA expression level of *FOS* after CPSF3 knockdown in wildtype and AFF4 MT cells. (D) The change of Pol II enrichment around the *FOS* gene after knockdown CPSF3 in WT and AFF4 MT cells analyzed by ChIP-qPCR. The *HEMO* gene acts as a negative control for ChIP-qPCR.

**Figure 7—figure supplement 1.**
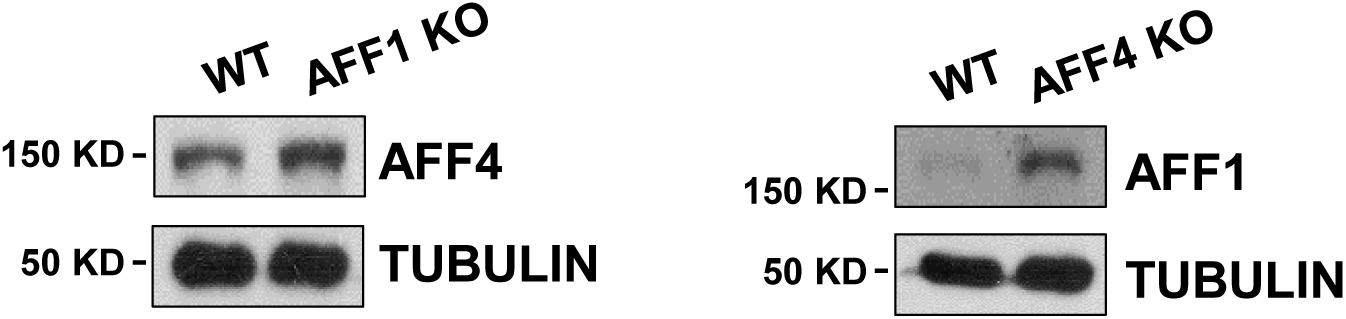
Significant increases of AFF4 occupancies at a subset of AFF1-depleted highly active genes. Western blot analyses showing the protein level change of AFF4 in AFF1 knockout and AFF1 in AFF4 knockout HCT116 cells. α-Tubulin was used as a loading control.

